# BAli-Phy version 3: Model-based co-estimation of Alignment and Phylogeny

**DOI:** 10.1101/2020.10.10.334003

**Authors:** Benjamin D. Redelings

## Abstract

**Summary:** We describe improvements to BAli-Phy, a Markov chain Monte Carlo (MCMC) program that jointly estimates phylogeny, alignment, and other parameters from unaligned sequence data. Version 3 is substantially faster for large trees, and implements covarion models, RNA stem models, and other new models. It implements ancestral state reconstruction, allows prior selection for all model parameters, and can also analyze multiple genes simultaneously.

**Availability:** Software is available for download at http://www.bali-phy.org. C_++_ source code is freely available on Github under the GPL2 License.

## Introduction

Accurate estimates of multiple sequence alignments (MSAs) are foundational to bioinformatics because MSAs are used as inputs when estimating gene trees, inferring positive selection, imputing ancestral sequences, estimating divergence times, and performing many other bioinformatic inferences. When the input alignments are not correct, errors may propagate downstream and undermine later inferences. For example, inferences of positive selection based on ClustalW alignments can lead to false-positive rates as high as 99% (Fletcher and Yang, 2010).

One approach to this problem is to use a model-based approach not only for downstream estimates such as ancestral sequences and gene trees, but also for the MSA. A realistic model of the MSA must place insertion-deletion (indel) events on specific branches of the phylogenetic tree, instead of merely placing gaps in a matrix (Loeytynoja and Goldman, 2008). Model-based inference then requires a probabilistic model of insertion-deletion events on each branch of the tree, as well as phylogenetic substitution likelihoods to score substitutions during alignment construction. A variety of programs implement model-based alignment, including Historian (Holmes, 2017), PRANK (Loeytynoja and Goldman, 2005), ProPIP (Maiolo *et al*., 2018). Model-based inference of alignments can increase alignment accuracy by up to 3-fold on simulated data sets (Redelings, 2014) when compared to the popular MSA construction tools MUSCLE (Edgar, 2004) and MAFFT (Katoh, *et al*., 2005).

However, simply using a better model for estimating the alignment fails to resolve a number of problems. First, constructing a single MSA estimate fails to account for alignment uncertainty. Second, model-based alignment requires an accurate tree estimate as input, but the accurate tree estimate requires an accurate alignment estimate as input. There is thus a “chicken and egg” problem of circular dependence between the alignment and the tree. These problems can both be solved by co-estimating the MSA and the phylogeny (Redelings and Suchard, 2005; Westesson *et al*., 2012; Arunapuram *et al*., 2013) in a Bayesian paradigm. This approach accounts for alignment uncertainty by providing a posterior distribution of alignments instead of just a single estimate of the alignment. Co-estimation solves the circular dependence problem because a high-quality internal estimate of the tree is constantly available.

Bayesian co-estimation of alignment and phylogeny is similar to Bayesian phylogeny estimation with a fixed alignment, as implemented in popular software such as MrBayes (Ronquist *et al*., 2012) or BEAST (Suchard *et al*., 2018). Both approaches are implemented using MCMC and output a collection of posterior samples instead of just a point estimate. Bayesian co-estimation extends fixed-alignment approaches by addition of (i) a probability model of insertion/deletion and (ii) MCMC transition kernels to propose new alignments. Each posterior sample then contains a phylogeny, an alignment, and numerical parameters. In addition, shared indels can be used to cluster taxa on the tree.

BAli-Phy is one of the leading software programs for performing Bayesian co-estimation of alignment, phylogeny, and other parameters (Redelings and Suchard, 2005; Redelings, 2014), along-side programs such as StatAlign (Arunapuram *et al*., 2013) and HandAlign (Westesson *et al*., 2012). Among these programs, BAli-Phy is currently unique in being able to analyze multiple partitions, specify priors for all variables, and log all sampled variables to a file. BAli-Phy can also fix or estimate the alignment in each partition independantly.

## Materials and Methods

### New features

#### Ancestral sequence reconstruction

Version 3 adds the ability to reconstruct ancestral DNA or amino-acid sequences. Unlike many programs including MrBayes, BAli-Phy reconstructs ancestral sequence with gaps, since it models indels. Each sampled alignment contains a “joint” reconstruction of ancestral sequences that corresponds to the co-sampled phylogeny. BAli-Phy can also summarize the sampled alignments to form a single “marginal” reconstruction of ancestral sequences. Here ancestral sequences are added to a consensus alignment and consensus tree in a way that averages over uncertainty in the topology and alignment.

#### Suitability for large trees

BAli-Phy version 3 is substantially faster when aligning large numbers of sequences (Figure 1). These improvements stem from the development of more efficient algorithms that are *O*(*n*) instead of *O*(*n*^2^) in the number of leaves *n*, including methods to avoid recomputing (i) the alignment matrix and (ii) cached conditional likelihoods when only part of the alignment changes. Version 3 also uses a branch length prior that does not pressure large trees to have huge total tree lengths, similar to the Dirichlet prior on branch lengths proposed by Rannala *et al*. (2012). Version 3 implements likelihood rescaling, so that likelihoods for large trees do not underflow.

**Fig. 1.**
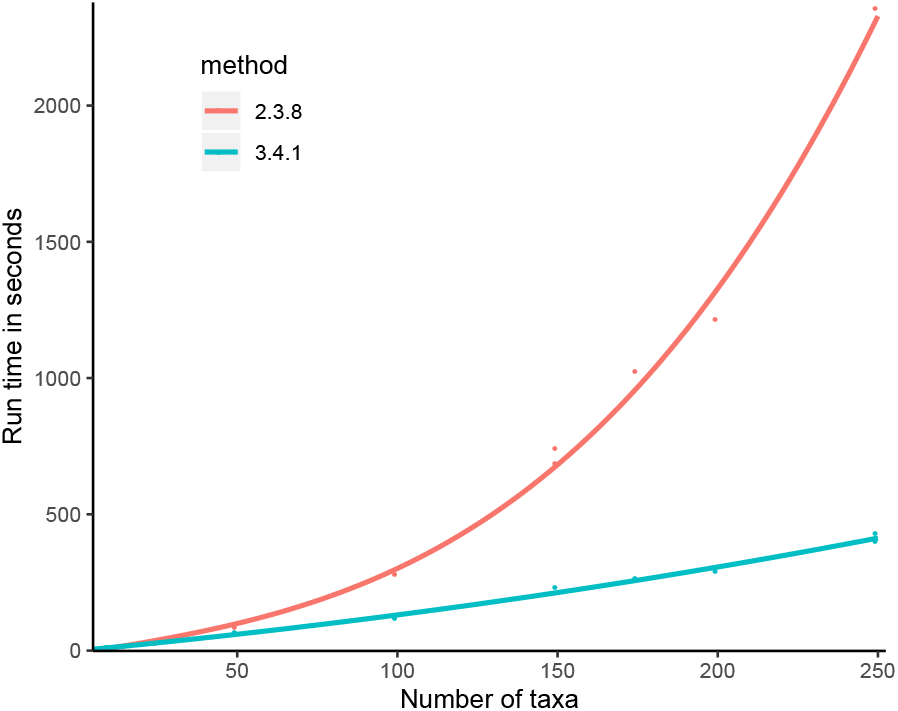
The time to run 10 iterations of MCMC versus the number of sequences used. The time increases quadatrically in version 2.3.8, but increases linearly in version 3.4.1.

#### MCMC improvements

Tree search has been enhanced to include subtree-prune-and-regraft (SPR) proposals, which allow larger moves within tree space than nearest-neighbor-interchange (NNI) proposals (Redelings and Suchard, 2007). When performing SPR, BAli-Phy uses an algorithm that considers attaching on every possible branch. This algorithm takes only *O*(*n*) time, where *n* is the number of leaves on the tree; it therefore improves exploration of topology space without paying a large computational penalty. BAli-Phy now uses slice-sampling to update numerical parameters and automatically tunes window sizes to achieve efficient mixing.

#### Explicit priors and fixed parameter values

BAli-Phy version 3 allows every evolutionary parameter to be given a fixed value or a prior. For example, to specify the *κ* parameter for the HKY model:

~~~
hky85 // default prior
hky85[kappa=2]
hky85[kappa^~^log_normal[log[2],0.25]
~~~

Default priors are now clearly reported to the user, so that it is always clear which priors have been used in an analysis. The ability to specify a fixed value for any parameter makes it possible to create unit tests for model likelihoods and to compare likelihood values with theoretical results and with other software programs.

The model language also allows a flexible stacking of models using the + operator, where `A+B` is equivalent to `B[submodel=A]`. For example:

~~~
lg08+f+Rates.gamma+inv
~~~

#### Codon and Triplet models

Version 3 adds the fMutSel and fMutSel0 codon models, in addition to the commonly used GY94 and MG94 codon models. Users can make use of the branch-site model and site-models such as M3 and M8a. Site-models can be applied to the MG94 and fMutSel models, in addition to the traditional GY94:

~~~
function[w,fMutSel[omega=w]]+m3
~~~

It is now possible to construct codon models in a flexible manner using model modifiers:

~~~
f81+x3+dNdS // mg94
gtr+x3+dNdS+mut_sel // fMutSel
~~~

#### RNA models

Version 3 adds multiple novel models for 16-state dinucleotide characters, which represent stems in RNA secondary structure. BAli-Phy cannot estimate secondary structure, so it relies on a pre-estimated alignment and secondary structure for RNA stems. BAli-Phy can be used to perform a traditional fixed-alignment analysis, placing stem columns in one partition and unpaired columns in another.

However, in some cases unpaired regions may be discarded because they are difficult to align. In such cases, it is possible to integrate out the alignment of unpaired regions by giving each unpaired region its own partition. Thus phylogenetic information in ambiguously-aligned unpaired regions can be retained without the need to trust any specific alignment. However, this scenario assumes that stem regions are sufficiently alignable to determine the boundary between stem and unpaired regions, even if the alignment within unpaired regions is uncertain.

#### “Covarion” and Markov-modulated models

We implement the Tuffley and Steel (1998), Huelsenbeck (2002), and Wang *et al*. (2007) models of heterotachy. These models allow rates of evolution to change over time *within* a site, instead of just between sites. For example the Tuffley-Steel approach models burstiness in evolution by creating an ON and OFF version of each DNA letter, where letters in the OFF state are invariant. The Huelsenbeck approach extends Tuffley-Steel to add Γ-distributed rate variation across sites, and the Wang *et al*. (2007) approach further extends this to allow changes in conservation over time.

We also implement Markov-modulated models, which allow switching between more general evolutionary models instead of just changing rates. For example, the Galtier (2001) model provides a rate *ν* for switching between different components in a mixture model. This allows switching between rates `(gtr+Rates.gamma+Covarion.gt01)`, different dN/dS values `(m3+Covarion.gt01)`, and more.

#### Visual mixing diagnostics for tree topologies

The `bp-analyze` program included with BAli-Phy generates an HTML report that summarizes MCMC runs. In addition to estimates of phylogeny, alignment, and parameters, this report includes several convergence diagnostics. These include a visual depiction of convergence in topology space. This method projects the Robinson-Foulds distances between sampled trees to 2 or 3 dimensions using principal components analysis (Hillis *et al*., 2005). The 3-dimensional projection can be visualized and rotated within a web browser.

#### Extensibility via packages

BAli-Phy version 3 adds the ability to extend the software via external packages. For example, the package BES makes use of the general MCMC machinery in BAli-Phy to infer self-fertilization rates from population samples under a coalescent model (Redelings *et al*., 2015). Packages are installable using the included `bali-phy-pkg` command, and are written using Haskell syntax.

### Analysis of real data

In order to assess run time on a real data set, we reanalyzed the ribosomal internal transcribed spacer (ITS) data set from Gaya *et al*. (2010). This analysis examined the 5.8S ribosomal RNA region and its two flanking ITS regions. The two ITS regions contain a large number of small indels and have a high evolutionary rate, whereas the 5.8S region has almost no indels and has a low evolutionary rate. We therefore divided the sequence data into 3 partitions, and fixed the alignment for 5.8S while allowing the ITS1 and ITS2 alignments to vary. To give an idea of the problem size, the ITS1, 5.8S, and ITS2 partitions have roughly 223, 156, and 172 columns, and the data set contains 68 individuals.

The two ITS partitions shared a linked `tn93+Rates.free[n=3]` model, but had unlinked `rs07` indel models. The 5.8S partition had a separate `tn93` model and no indel model. We ran four unheated chains under version 3.5 for 24 hours on a heterogenous cluster. We performed the same analysis on version 2.3.8 in the same environment for comparison. Command files are included in the supplement.

## Results

The BAli-Phy 3.5 analysis reached ASDSF=0.007 and MSDSF=0.076 in 24 hours, completing 11261, 12669, 10591, 10672 iterations. A value of 0.01 for the ASDSF is commonly taken to indicate convergence and sufficient sampling. The BAli-Phy 2.3.8 analysis reached ASDSF=0.017 and MSDSF=0.186 during the same time period, completing 5050, 4823, 5665, 5688 iterations. (Note that BAli-Phy iteration numbers are not comparable to MCMC software like MrBayes because BAli-Phy does a lot more work in each iteration, updating the alignment, tree, and each numerical parameter.)

As described in Gaya *et al*. (2010), co-estimating the alignment and tree results in greater support for many branches. This is because indels that occurred on those branches can be used to group taxa. Thus, substitutions are no longer the sole basis for estimating support. Figure 2 depicts the consensus alignment for ITS1 and the majority consensus tree.

**Fig. 2.**
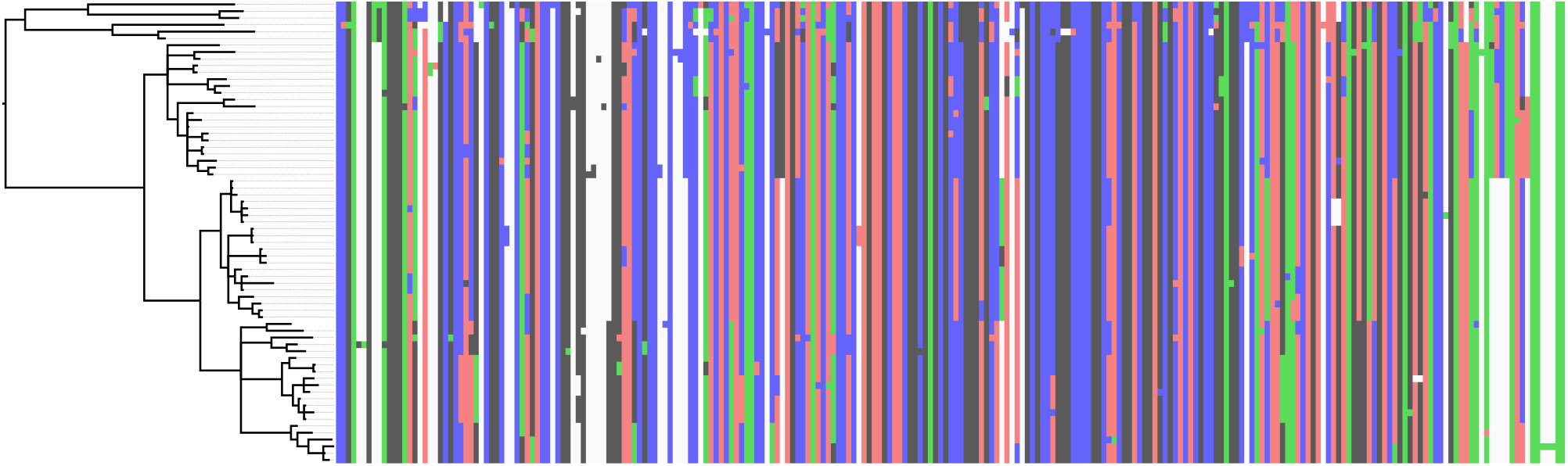
The phylogeny estimate (majority consensus) and alignment estimate (maximum posterior decoding) for ITS1. Many of the branches are supported by shared indels.

## Conclusion

These results indicate that BAli-Phy version 3 can perform a complete analysis of this data set in 24 hours on modern hardware. This is substantially faster than the 15 days taken by a similar analysis with version 2.1 in 2009 (Gaya *et al*., 2010). This speed increase is partly the result of faster computing equipment. However, our results indicate that version 3 is roughly twice as fast as version 2.3 on the same hardware for a dataset of 68 taxa. Run time remains *O*(*L*^2^) in sequence length *L* for each partition.

## Acknowledgements

B.D.R. was partially supported by NSF-DBI-1759838 (“Collaborative Research: ABI Development: Cultivating a sustainable Open Tree of Life”) and by NSF-DEB-1256993 (“Integrating Fossil Data into Likelihood-based Phylogenetic Analyses with Trilobites as a Model System”).

